# A Functional Schizophrenia-associated genetic variant near the *TSNARE1* and *ADGRB1* genes

**DOI:** 10.1101/2023.12.18.570831

**Authors:** Marah H. Wahbeh, Rachel J. Boyd, Christian Yovo, Bailey Rike, Andrew S. McCallion, Dimitrios Avramopoulos

**Affiliations:** McKusick-Nathans Department of Genetic Medicine, Johns Hopkins University School of Medicine, Baltimore, MD 21205, USA; Department of Medicine, Johns Hopkins University School of Medicine, Baltimore, MD 21205, USA

**Keywords:** gene regulation, genome editing, CRISPR, genetic association, schizophrenia, psychosis, *TSNARE1*, *ADGRB1*, synapse, neurons, axons, neurodevelopment, zebrafish, induced pluripotent stem cells

## Abstract

Recent collaborative genome wide association studies (GWAS) have identified >200 independent loci contributing to risk for schizophrenia (SCZ). The genes closest to these loci have diverse functions, supporting the potential involvement of multiple relevant biological processes; yet there is no direct evidence that individual variants are functional or directly linked to specific genes. Nevertheless, overlap with certain epigenetic marks suggest that most GWAS-implicated variants are regulatory. Based on the strength of association with SCZ and the presence of regulatory epigenetic marks, we chose one such variant near *TSNARE1* and *ADGRB1*, rs4129585, to test for functional potential and assay differences that may drive the pathogenicity of the risk allele. We observed that the variant-containing sequence drives reporter expression in relevant neuronal populations in zebrafish. Next, we introduced each allele into human induced pluripotent cells and differentiated 4 isogenic clones homozygous for the risk allele and 5 clones homozygous for the non-risk allele into neural precursor cells. Employing RNA-seq, we found that the two alleles yield significant transcriptional differences in the expression of 109 genes at FDR <0.05 and 259 genes at FDR <0.1. We demonstrate that these genes are highly interconnected in pathways enriched for synaptic proteins, axon guidance, and regulation of synapse assembly. Exploration of genes near rs4129585 suggests that this variant does not regulate *TSNARE1* transcripts, as previously thought, but may regulate the neighboring *ADGRB1*, a regulator of synaptogenesis. Our results suggest that rs4129585 is a functional common variant that functions in specific pathways likely involved in SCZ risk.

## 1. INTRODUCTION

Schizophrenia (SCZ) is a chronic, neuropsychiatric disorder generally characterized by a set of symptoms that include delusions and hallucinations; disorganized speech and behavior; and cognitive impairments, such as mental, emotional, and social deficits^1^. SCZ is estimated to impact over 24 million individuals worldwide, with particularly high prevalence, incidence, and burden in the United States^2^. In addition to the mental, social, occupational, and educational duress associated with SCZ, affected individuals have a reduced life expectancy by about 12-15 years relative to the general population^3–5^.

While the genetic contribution to SCZ risk is high, with a heritability of 64%-80% ^6–8^, patients exhibit variable phenotypes and complex inheritance patterns. This is indicative of an environmental contribution to disease risk mediated by gene-by-environment interactions^7, 9^. Current treatment strategies for SCZ include the administration of antipsychotic drugs. However, many clinical trials for antipsychotics have failed or yielded negative-to-moderate effects in SCZ patients, who exhibit variable responses to treatment depending on age of onset, sex, geographic location, and other factors^10^. This clinical heterogeneity is thought to reflect the contribution of varying underlying biological processes among patients^11^. Therefore, in order to develop more effective or targeted therapeutics, it is imperative that we understand the genetic architecture of this disorder.

In recent years, collaborative large genome wide association studies (GWAS) have robustly identified 287 independent loci contributing to the risk for SCZ^12^. While these discoveries are valuable, one of the primary limitations of a GWAS is that it cannot resolve the lead single nucleotide variant (SNV) driving the risk association, from other variants in linkage disequilibrium (LD). It is also increasingly acknowledged that most variants identified by GWAS are located in non-coding regions of the genome, and ∼40% of the time, the haplotype blocks that contain these lead SNVs do not include coding exons^13–15^. In fact, a growing body of evidence, including expression Quantitative Trait Loci (eQTL) studies, suggest that most variants in LD with lead GWAS SNVs are regulatory in nature^15–19^ and concentrate in regulatory DNA marked by deoxyribonuclease I (DNase I) hypersensitive sites (DHSs)^20^. Nevertheless, even when variants are eQTLs for specific genes, it is often not just one gene that they regulate, and the LD conundrum remains. As a result, genes are often implicated because of their proximity to SCZ-associated variants and known involvement in synaptic biology but are seldom accompanied by direct functional evidence.

While identifying reliable genetic associations is a significant first step towards understanding the etiology of SCZ, functional studies linking specific variants to the regulation of specific genes and biological processes is an important next step towards unlocking their translational potential. Understanding how SCZ associated variants are involved in regulatory activities will promote the generation of innovative disease treatments and prophylactics. Technological advancements in cell engineering and genome editing using CRISPR/Cas9^21^ – and the ability to generate human induced pluripotent stem cells (hiPSCs) that can be differentiated to a variety of cell types – have created a new toolkit for modeling human diseases^22^. Neurons that carry variants implicated in disease risk can now be generated and compared to neurons that carry non-risk alleles at the same position, on an otherwise isogenic background. This strategy can be used to not only show the functionality of non-coding, potentially regulatory variants, but also to observe its consequences in a disease-relevant context. One of the most informative ways to characterize and phenotype such allelic differences is to study the transcriptome, in which allele-specific effects may be identified that impact the target gene, but also point to disrupted transcriptional programs and their pathways.

We employ this strategy in the work we present here, where we test the functionality of one such non-coding, SCZ-associated, SNV based on the strength of its association to SCZ and the presence of open chromatin marks in a disease-relevant, neuronal population. We first employ a well-established transgenic zebrafish reporter assay^23–25^ to confirm the regulatory potential of the sequence within which this variant resides. We then use precise genome editing to generate isogenic hiPSC-derived neural precursor cells (NPCs) that possess the risk allele and non-risk allele and compare the transcriptional differences between NPCs with the SCZ risk allele versus the non-risk allele. Our results support the functionality of this SCZ risk variant, identify transcriptional changes associated with possession of this variant, and unearth additional downstream effects within SCZ-related pathways in the transcriptome.

## 2. MATERIAL AND METHODS

### 2.1. Variant Prioritization

We prioritized characterization of variant rs4129585 based on the information available at initiation of the project. The second version of SCZ GWAS data was available at the time from the SCZ working group of the psychiatric genetics consortium (PGC) (https://pgc.unc.edu/for-researchers/working-groups/schizophrenia-working-group/)^26^, which indicated that rs4129585 was the top-ranking SNV at that locus near the *TSNARE1* and *ADGRB1* genes (*p*= 2.03×10^-13^, risk allele = C). This variant was reported by the PICS fine-mapping algorithm (pics2.ucsf.edu) to be the most likely driver of SCZ association at that locus. Using the UCSC genome browser, we found that rs4129585 overlaps with a chromatin immunoprecipitation signal from restrictive element-1 silencing transcription factor / neuron-restrictive silencing factor (REST/NRSF) and was shown to drive reporter gene expression in K562 cells (p=2.4 x10^-4^) in a massively parallel reporter assay (MPRA) we recently reported^27^. Of note, another nearby SNV, rs13262595, is in very high LD with rs4129585 (LD, r^2^=0.996 from https://ldlink.nih.gov/), which was also strongly associated with SCZ risk. At the time, this SNV showed slightly weaker association (*p=* 2.85×10^-13^, risk allele = G); however, in most recent PGC data^12^, rs13262595 had slightly stronger signal (*p*= 5.109×10^-18^) compared to rs4129585 (*p*= 7.062×10^-18^). It also overlaps with many chromatin marks, making it another good candidate to drive SCZ risk at this locus. This SNV was not found to drive reporter gene expression in our published MPRA^27^. Nevertheless, rs13262595 remains a target of our future investigations.

### 2.2. Functional Assessment using Zebrafish Transgenesis

To evaluate the regulatory potential of rs4129585 and its surrounding sequence, we employed Tol2 transposon-mediated transgenesis according to our standard protocol^23–25^, with minor modifications. Briefly, primers were designed to amplify a human non-coding sequence containing rs4129585 (**Table S1**). The boundaries of this sequence (hg38 chr8:142231443-142231752) were chosen to include a ChIP-seq peak surrounding rs4129585, as identified by the UCSC Genome Browser hg38 track “Transcription Factor ChIP-seq Clusters (340 factors, 129 cell types) from ENCODE 3,” suggestive of restrictive element-1 silencing transcription factor (REST)/neuron-restrictive silencing factor (NRSF) transcription factor (TF) occupancy.

Next, we ligated 5′ *att*B sequences to each primer set and amplified our chosen sequence by PCR. Instead of gel extraction, leftover PCR product was purified using the DNA Clean & Concentrator™-5 kit (Zymo Research #D4067). The *attB* PCR product was cloned into an empty pDONR™221 vector, and the resulting construct underwent recombination with a pGW_cfosEGFP destination vector, such that the conserved non-coding sequence was placed upstream of a *cfos* minimal promoter that drives EGFP expression and flanked by Tol2 transposons that promote efficient genomic integration of the reporter construct. Purified recombinant DNA was injected into 1–2-cell zebrafish embryos, as previously described^23^. Injections were performed in zebrafish embryos of the strain AB, which were obtained from the Zebrafish International Resource Center (http://zfin.org; ZFIN ID: ZDB-GENO-960809-7) and raised in our facility. Embryos were screened for EGFP expression at 4- and 6-days post fertilization (d.p.f.) using a Nikon AZ100 Multizoom fluorescence microscope, and the embryos with the strongest EGFP signal were selected to be raised to sexual maturity.

After 3 months, ≥5 sexually mature adults were crossed with wild-type AB fish and F_1_ embryos were screened for EFGP to confirm germline transmission of the transposon-mediated insertion, again using a Nikon AZ100 Multizoom fluorescence microscope at 6 d.p.f. To ensure that position effects of individual transgene insertions did not confound our results^23^, F_1_ embryos were screened from each of 5 independent parental clutches, and ≥3 representative F_1_ founder fish with consistent EGFP expression were selected from each clutch and raised to sexual maturity. At 3 months of age, F_1_ founders were again crossed with wild-type AB fish. The resulting F_2_ embryos were collected and raised in embryo media (E3) that contained 200 μM 1-phenyl 2-thiourea (PTU; Sigma Aldrich #P7629) to inhibit pigmentation^28^. This E3/PTU media was replaced daily.

At 6.d.p.f., ethyl-3-aminobenzoate methanesulfonate salt (Tricaine/MS-222; Sigma Aldrich #E10521) was prepared at a concentration of 4 mg/mL, buffered to pH7 using 1M Tris (pH 9; 4% v/v), and added to E3 medium to anaesthetize zebrafish larvae during the imaging process. Imaging was performed using the GFP filter and auto exposure settings on a Nikon AZ100 Multizoom fluorescence microscope, and GFP expression patterns among founder zebrafish were analyzed (**Table S2**), after which zebrafish larvae were transferred to a new dish containing fresh E3 media.

### 2.3. Cell line used for CRISPR editing and cell culture

The iPSC line MH0180966 used for CRISPR/Cas9 editing and further experiments was derived from a female individual of European descent without a psychiatric diagnosis and provided to us by the stem cell repository at Rutgers University (NIMH/RUCDR). We received the line at passage number 12. The iPSC line was treated as we have outlined previously^21, 29^. Briefly, the cells were grown on culture plates coated with 5μg/mL laminin (ThermoFisher BioLamina #LN521) and incubated in a humidified 5% CO_2_ incubator at 37°C for ≥2 hours. Once seeded, the cells were maintained in StemFlex media with supplement (ThermoFisher Gibco #A3349401) and 1% penicillin–streptomycin (Penn/Strep; Gibco # 15140122) in a humidified 5% CO_2_ incubator at 37°C. When thawing, passaging, or transfecting, 10μM Y-27632 Rho-kinase inhibitor (ROCKi; Tocris #1254) was added to culture media. Cells at 70-80% confluence were passaged using 1X Phosphate-buffered saline (PBS) and accutase (MilliporeSigma #A6964) every ∼4-5 days. Cell stocks were first frozen at −80°C in StemFlex media with 10% Dimethylsulfoxide (DMSO) for 24h before storage in liquid nitrogen. iPSC genotyping was done by PCR and Sanger sequencing using primers listed in the Supplementary Information file of Feuer & Wabeh, 2022 (see Online Methods; *TSNARE1* experiment)^21^.

### 2.4. Generating Allele-Specific iPSC Lines

To induce target sequence deletion and subsequent re-insertion of the rs4129585 non-risk (A) and risk (C) alleles, we employed our pipeline for scarless genome editing, CRISPR Del/Rei^21^. Detailed methods, transfection and selection details, and summary of off-target analyses; as well as sequences for primers, single guide RNAs (sgRNAs), and single stranded oligodeoxynucleotide (ssODN), can be found in the Supplementary Information file of Feuer & Wahbeh *et al*., 2022 (see Online Methods; *TSNARE1* experiment)^21^. In brief, sgRNA oligos were designed using CRISPOR (http://crispor.tefor.net/crispor.py) to flank rs4129585 and induce a deletion between GRCh37/hg19 chr8:142231553-142231597. Restriction enzyme BbsI sticky ends were added to single-stranded DNA oligos before the oligos were annealed and cloned in pairs into pDG459 plasmids containing a puromycin resistance gene. pDG459 plasmids were transfected into iPSCs using lipofectamine-stem and successful transfection was selected for using puromycin^21^. Cells were screened for desired edits and candidate off-target effects^21^ at 4 genomic locations that showed 3 or fewer mismatches with the gRNAs and were in coding sequences or overlapped with DNAse hypersensitivity sites from the encode project (www.encodeproject.org; file hg38_wgEncodeRegDnaseClustered.txt). Single cell clones with the appropriate deletion and no off-target events were chosen for expansion. Next, a synthetic sgRNA (syn-sgRNA) and an ssODN repair template were designed to reinsert either the non-risk allele or risk allele at the appropriate genomic position. As with the deletion step, syn-sgRNAs were cloned into a pX459 plasmid with a puromycin resistance gene, transfected into iPSCs, and edited cells were selected for using puromycin, before iPSCs were again screened for appropriate reinsertion and lack of off target effects at 8 candidate genomic locations, as above ^21^.

### 2.5 Differentiation into Neural Precursor Cells

Each iPSC clone was differentiated into Neural Progenitor Cells (NPCs) through directed differentiation of embryoid bodies^30^ (EB) with SMAD, as previously described^31, 32^ with minor modifications. In brief, iPSCs were first plated at a density of ∼300 k/well in 3 wells of 6-well laminin-coated plate. Differentiation was initiated 2 days later by passaging cells at a density of 5K cells/well in a round-bottom, untreated 96-well plate (Corning #3788) in 50ul/well of MTeSR media (Thermo #85850) and Rocki (Y-27632 dihydrochloride; BD Biosciences # 562822). Directly after plating, the plate was spun at 1,000 RPM for 3-4 minutes to condense the cells and facilitate EB formation. The following day, (DIV0) the presence of EBs were confirmed and 50uL of Neural Induction Media (NIM) was added to each well, composed of DMEM/F12 Media (Thermo #11320033) with 2% B27 supplement without vitamin A (Thermo #12587010), 1% GlutaMAX (Thermo # 35050061), 1% N2 (Thermo #17502048), 1% MEM NEAA (Non-Essential Amino Acids; Thermo #11140050), and 1:100 BME (Beta-mercaptoethanol; Thermo #21985023) with freshly added SMAD Inhibitors SB431542 (Reprocell #NC1388573) at 20uM and LDN193189 (Peprotech #1062443 -1mg) at 200nM. From DIV1-DIV3, 50ul of NIM was added to each well with half the concentration SMAD Inhibitors as day 1 (SB431542 at 10uM, LDN193189 at 100nM) until reaching 200ul/well.

From DIV4 onwards, 100ul of NIM with SMAD Inhibitors was exchanged daily until EB maturity (DIV6 - DIV9). EBs were passaged into 3 wells of a 6-well plate that was coated in 1:100 matrigel (Corning # 47743-720) for 1 hour at 37°C and contained 2ml of NIM/well with SMAD Inhibitors. EBs were then carefully removed from the 96-well plate into 1-2 wells of an uncoated 6-well plate with warm DMEM/F12, before being transferred to the coated plate containing NIM. The EBs adhered to the Matrigel and were fed daily from DIV7-DIV13 with NIM, SMAD inhibitors, and 10ng/mL FGF2 (Miltenyi #130-093-838) until robust rosettes formed. Once formed, rosettes were manually picked and transferred into 3 wells of a 6-well plate coated in 1:100 matrigel for 1 hour at 37°C and containing 2ml of NIM/well with SMAD inhibitors. The following day onwards, cells were fed with NIM, SMAD inhibitors, and FGF2 until robust rosettes formed again. The rosettes were again lifted manually, and Accutase was added to separate the cells before they were plated into NPC media containing half DMEM/F12 and half Neurobasal Media (Thermo #21103049) with 2% B27 supplement without vitamin A, 1% GlutaMAX, 1% N2, 1% MEM NEAA and 1:100 BME. FGF2, 10ng/ml EGF (Peprotech #100-47-10ug) and Rocki were also added. Cells at this stage were fed with this NPC media without Rocki every 2 days until ∼70% confluent. After 2 passages, the cells were characterized by the presence of NPC markers.

### 2.6 Characterization of NPCs

To confirm the identity and quality of NPCs, we visualized the expression of NPC markers and lack of expression of iPSC markers by immunocytochemistry staining. 3-4 days before staining, a 4-well chamber slide (Thermo #62407-294) was coated with 1:100 matrigel in DMEM/F12 for 1 hour (300uL per well) and ∼150k NPCs were plated into each well. Slides were fed once every 2 days with NPC media until the cells were ∼60% confluent. NPC media was then removed, and the cells were washed with cold 1XPBS and fixed using 4% PFA (Alfa Aesar #J61899). Cells were blocked for 1 hour using a blocking buffer composed of 1XPBS, 5% Goat Serum, and 0.3% Triton X-100 (Sigma #93443-100mL), after being washed three times with 1X PBS for 5 minutes each time. Following blocking, cells were incubated in antibody solution made up of 1XPBS, 1% BSA, and 0.3% Triton X-100, as well as different combinations of 1:200 Nestin antibody, 1:200 PAX6 antibody, 1:300 TRA-160 antibody, 1:300 SOX2 antibody, and 1:200 Nanog antibody at 4°C overnight. Slides were washed three times with cold 1X PBS for 5 minutes each time before adding antibody solution with 1:400 488 Goat α-Mouse secondary antibody and 1:400 565 Goat α -Rabbit secondary antibody. Incubation lasted 1h at 37°C before washing three times with cold 1X PBS again before mounting with 25uL of the prolong mounting media with DAPI (Invitrogen #P36931) to each well. Slides were imaged with an Axio Observer fluorescent microscope. Next, we performed RT-qPCR in some of the samples by generating cDNA from 1ug of RNA from each clone before and after differentiation (iPSCs and NPCs) using the Superscript III kit, following the manufacturer’s protocol (ThermoFisher # 18080400). cDNA was diluted 1:10 in water for RT-qPCR. We used iTaq and custom primers (**Table S3)** following the manufacturer’s protocol (BioRad # 1725120). We used the CFX Connect System with denaturation at 95°C for 20 sec, amplification for 40 cycles, Denaturation at 95°C for 1 sec, annealing at 60°C for 30, and Melt Curve Analysis at 65-95°C. Normalized relative quantification and error propagation was calculated and analyzed as proposed by Taylor *et al.* (2019)^33^, with results normalized to *ACTB* and *GAPHD*. A two-sample t-test was performed using R function “t.test”, (alternative = “two.sided”) to test for significant differences in gene expression.

### 2.6. RNA-sequencing

Total RNA was isolated from 1 confluent well of NPCs/line from a 6 well plate using the Zymo Quick-RNA™ Miniprep Kit (#R1054) following the manufacturer’s protocol. RNA was submitted to Novogene (novogene.com) for 150-bp paired-end RNA-sequencing. All libraries passed Novogene’s post-sequencing quality control, except REF_5, which was excluded from analysis. We received an average of 51.1 million reads per sample with a maximum of 74.9 and a minimum of 42.5 million reads. Reads were aligned to the human reference genome GRCh38 using Hisat2 version 2.1.0^34^ and corresponding BAM files were generated using Samtools 1.9^35^. Stringtie 2.0.3^36^ was used to assemble and estimate the abundance of transcripts based on GRCh38 human gene annotations^37^. Bioconductor package tximport was used to compute raw counts by reversing the coverage formula used by Stringtie with the input of read length^36^. The output was then imported to the Bioconductor package, DESeq2, for differential gene expression analysis^38^. Benjamini-Hochberg adjustment^39^ was used to calculate adjusted p-values (padj). Before expression analysis, principal component (PC) analysis (**Figure S6; Figure S7**) was performed on the adjusted read counts and the first PC was included in further analysis as a covariate.

### 2.7 Bioinformatics Analysis

To explore which genes were differentially expressed (DE) between edited and unedited iPSC-derived neurons (rs4129585 non-risk and risk alleles), we used the Panther bioinformatics platform^40^ (https://www.pantherdb.org/), which reports enrichments across the Gene ontology (GO) resource terms (https://geneontology.org/) corrected for multiple testing. We used all genes with detectable transcripts in our dataset as a background. To look for interactions between the DE genes, we used the STRING database^41, 42^ (string-db.org). As suggested by the curators of the STRING database website, we used the medium-level confidence score (0.4) for interactions and did not include any additional level of interactors, to ensure the validity of the observed network enrichment p-values. In addition to enrichment for interactions, STRING reports significant functional enrichments, similar to Panther but tested only among interacting DE genes, thus pointing to the function of the identified network.

## 3. RESULTS

### 3.1. A transcription factor binding site containing rs4129585 drives reporter gene expression in transgenic zebrafish neuroanatomy

To test rs4129585 and the surrounding sequence for regulatory function, we utilized a transgenic zebrafish reporter assay in which the presence of an upstream regulatory element is required for a minimal promoter to induce GFP reporter gene expression. The SCZ-associated variant rs4129585 lies within a ChIP-seq REST/NRSF TF binding site (TFBS), and the entire sequence representing the ChIP-seq peak was integrated into the zebrafish genome upstream from a *cfos* minimal promoter/GFP reporter construct. REST is an epigenetic remodeler that is widely expressed during human embryogenesis and late-stage neuronal differentiation. REST acts as a transcriptional repressor of neural-specific genes involved in synaptogenesis, axonal pathfinding, synaptic plasticity, and structural remodeling in humans^43, 44^; therefore, it is unsurprising that this sequence drives GFP expression in the zebrafish brain (**Figure 1)**.

**Figure 1.**
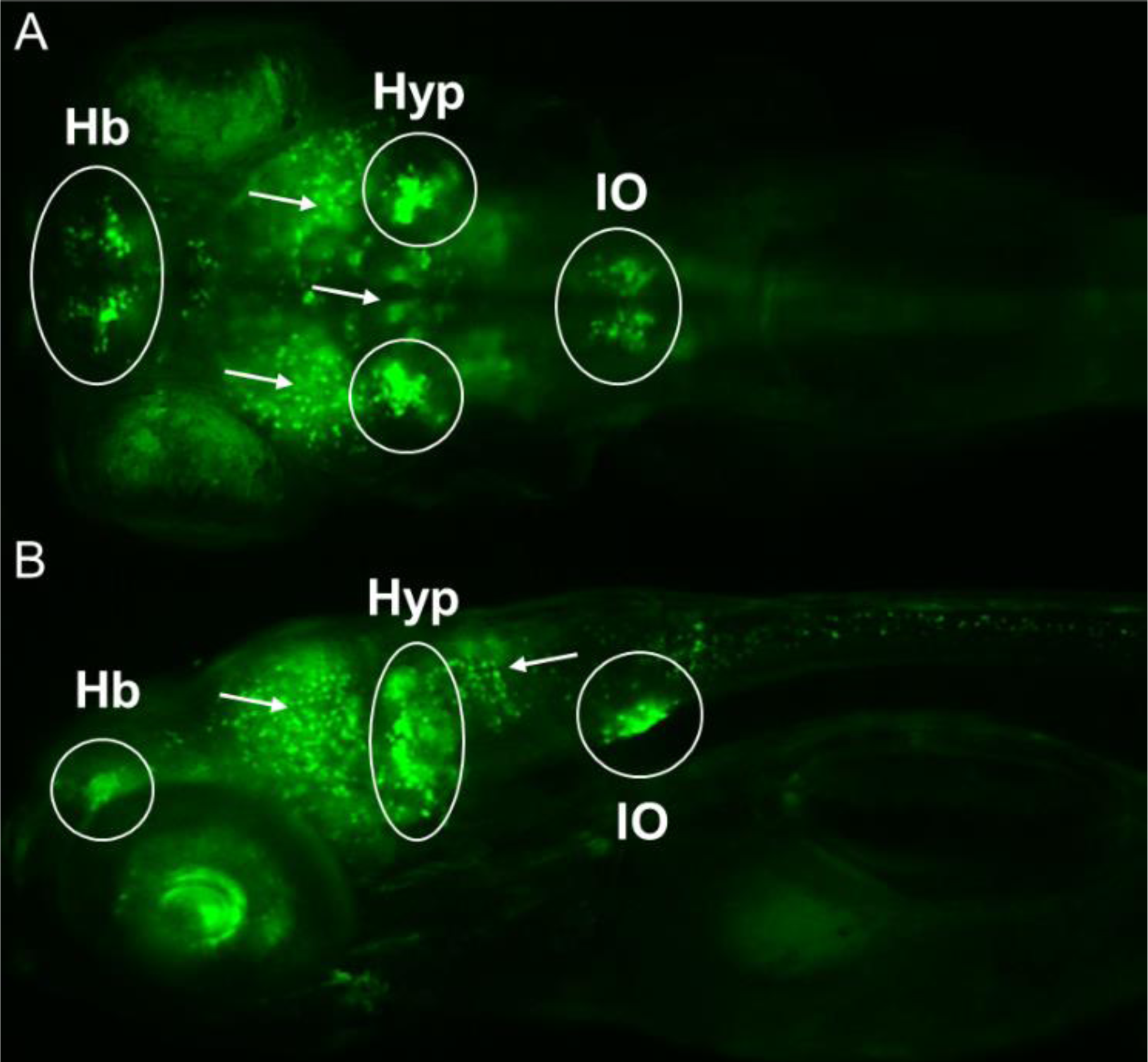
A noncoding REST/NRSF TF binding sequence containing rs4129585 drives reporter expression in F_2_ zebrafish embryos 6 d.p.f. **(A)** Dorsal and **(B)** lateral views of a representative zebrafish embryo with GFP expression patterns in the habenula (Hb), hypothalamus (Hyp), and inferior olive (IO), as well as glutamatergic neurotransmitters (arrows).

Representative GFP expression pattern from F_2_ zebrafish embryos suggest that this sequence is functional in three major neuroanatomical regions (**Figure 1 & Figure S1**): (1) the habenula, which is involved in aversive social responses in zebrafish^45^, including anxiety^46^, fear^47^, and aggression^48^; (2) the caudal hypothalamus, which is involved in appetite control^49^, but in which dopaminergic (DA) neurons are also involved in audio-sensory functions, including locomotion and sensation of acoustic/vibrational stimuli^50^; and (3) the inferior olive, which receives DA projections from the caudal hypothalamus^50^, but also provides climbing fibres to Purkinje cells of the cerebellar cortex that are required for motor learning^51, 52^. The more widespread GPF signal in these fish is consistent with expression in glutamatergic neurons, as well as a few regions with GABAergic neurons (**Figure 1**). These findings suggest that the SCZ-associated variant, rs4129585, likely lies within a TFBS that elicits its regulatory function in glutamatergic neurons and regions of the brain that have been implicated in neurodevelopment and SCZ pathology^53–62^.

### 3.2. iPSC-derived NPCs exhibit transcriptomic changes associated with a SCZ risk allele

With evidence that rs4129585 lies within a functional regulatory element implicated in neurobiology, we obtained human-derived iPSCs, used CRISPR-mediated genome editing to engineer the rs4129585 risk and non-risk alleles into otherwise isogenic cell lines, and differentiated these lines into NPCs. NPCs were selected as the differentiation endpoint because they are at an early stage in neuronal differentiation, and SCZ is considered a neurodevelopmental disorder^59, 60, 63, 64^. Immunostaining and RT-qPCR with NPC- and iPSC-specific markers showed the former to be upregulated and the later downregulated (**Figure S2, Figure S3, Figure S4, Figure S5**); thus, confirming successful differentiation from iPSCs to NPCs. We then performed RNA-seq to identify the transcriptional changes elicited by the altered variants. PC analysis of the RNAseq data identified one reference clone (REF_6) to be an outlier across PC1 (**Figure S6**). NPC and iPSC marker gene expression (**Table S4**) suggest that this cell line might not be as far along in the differentiation process. To correct for this difference going forward, we used PC1 as a covariate in the analysis. Differential expression analysis revealed differences in the expression of 109 genes at FDR <0.05 and 259 genes at FDR <0.1. Of these, 116 were upregulated and 143 were downregulated (**Table S5** shows the complete results at the gene level, while **Table S6** shows the results at the transcript level). We then proceeded to test these genes for functional enrichments using the Panther Bioinformatics database^40^, using the FDR <0.1 threshold to obtain higher statistical power. This revealed strong enrichments in many functional categories with direct relevance to schizophrenia, such as neuron development, differentiation, axonogenesis, and axon guidance, among many others (**Figure 2**).

**Figure 2.**
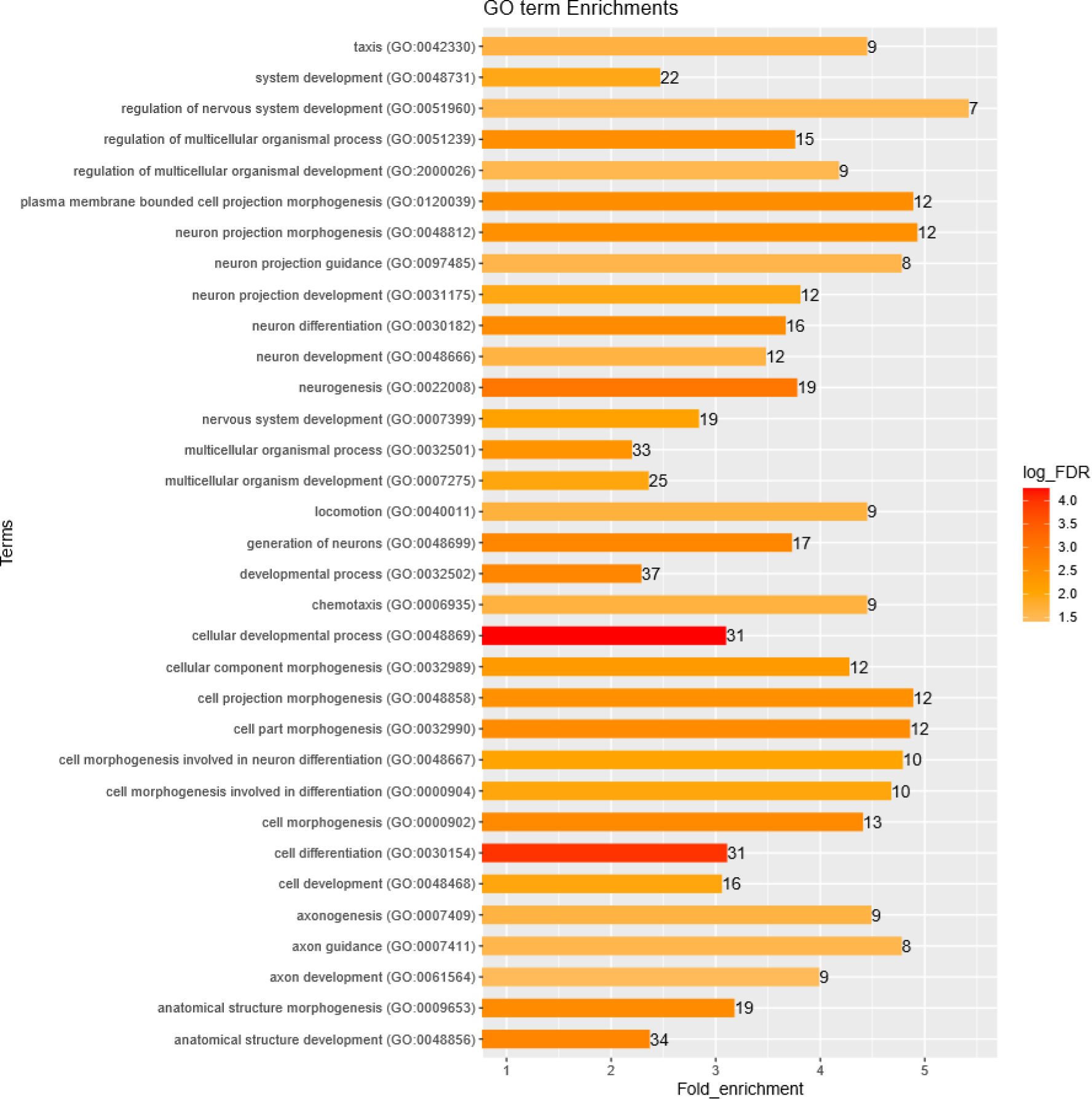
GO terms for functional enrichments among DE genes reported by PANTHER. Bar color indicates level of statistical significance (log_FDR; false discovery rate), numbers indicate the number of genes enriched in the pathway, and bar length indicates the fold enrichment of the genes observed in our uploaded list over the expected representation of genes in each GO category.

We next used the STRING database of protein-protein interactions^41, 42^ to test whether the protein products of the dysregulated genes are functionally related as evidenced by reported interactions between them, an indication of validity of the results which also allowed further functional exploration. This analysis showed that these genes are highly interconnected (STRING enrichment-*p=*1.2×10^-11^) with 1.66-fold more interconnections than expected by chance for the same number of proteins. Further, functional enrichment analysis for the network proteins as performed by STRING showed many of the same enrichments observed with Panther Bioinformatics, now including synapse organization and assembly, (FDR<0.02), central nervous system development, regulation of synapse assembly and axon guidance. **Figure 3** shows all enrichments and **Figure 4** shows as an example the 11 axon guidance genes in the STRING network, representing a more than 4-fold enrichment from the expected.

**Figure 3.**
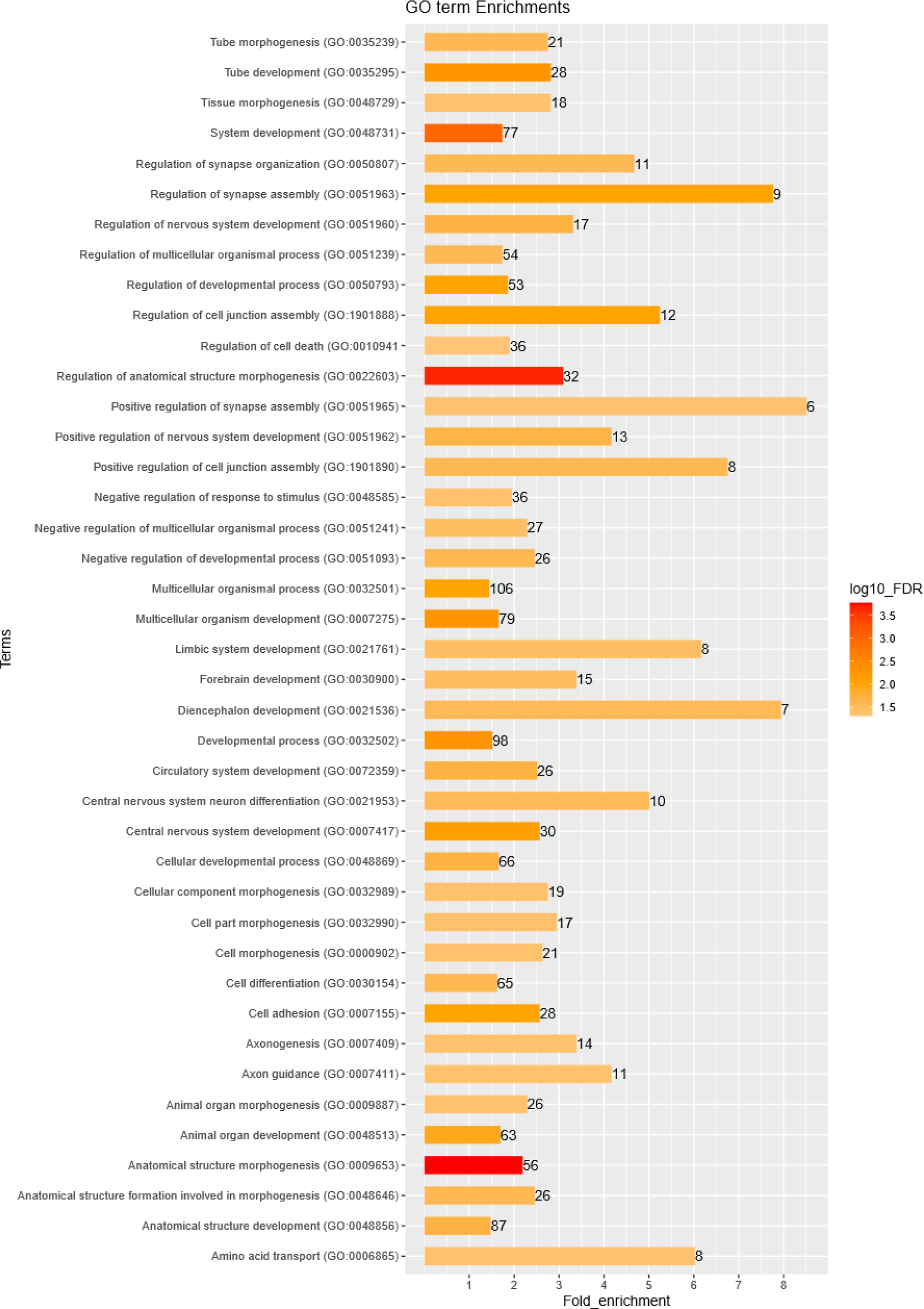
GO terms for functional enrichment within the STRING network of interacting proteins identified from our DR genes. Bar color indicates level of statistical significance, numbers indicate the number of genes enriched in the network, and bar length indicates the fold enrichment of the genes observed in our uploaded list over the expected representation of genes in each GO category.

**Figure 4.**
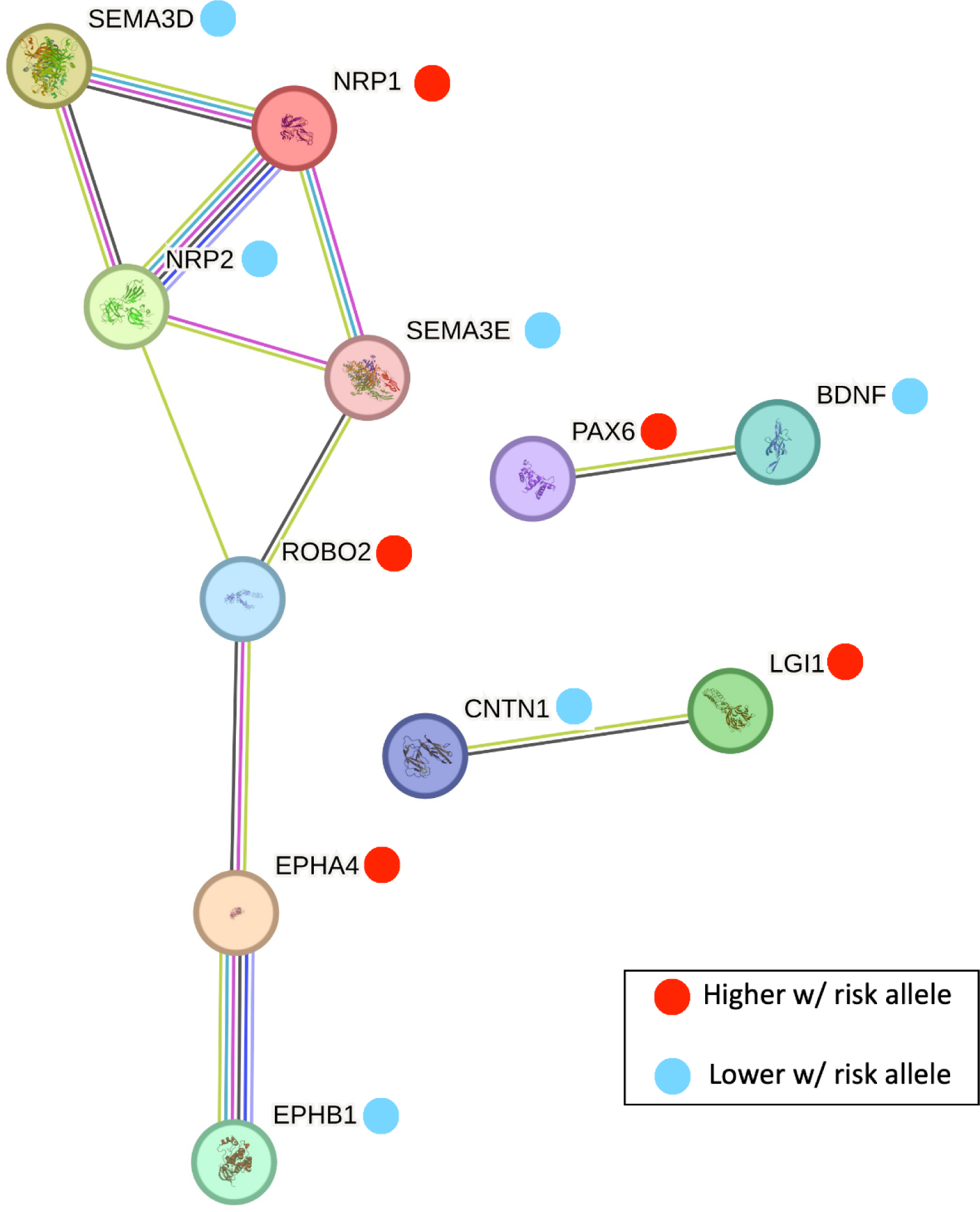
The 11 axon guidance genes in the STRING network of DE genes. Blue dots indicate proteins in the network whose genes are downregulated in the NPCs containing the risk allele, and red dots indicate proteins in the network whose genes are upregulated in the NPCs containing the risk allele. Image generated by the STRING website.

Results from the Genotype-Tissue Expression (GTEx) project (www.gtexportal.org/home/) show that rs4129585 is an eQTL for its’ two nearest genes, *TSNARE1* and *ADGRB1,* regulating both the genes and specific transcripts’ abundances in a variety of tissues and at varying directions (including cerebellum but not brain). We looked at our results for the effects of the variant on these two genes and found no effect on *TSNARE1* and a suggestive effect on *ADGRB1*(*p*=0.06), with the risk allele increasing expression in our induced NPCs. Interestingly, *ADGRB1*, is a regulator of synaptogenesis^65^. None of the transcripts of *TSNARE1* (9 present in our data) showed significant differences. Of the 5 *ADGRB1* transcripts in our data, two showed significant differences (ENST00000517894, nominal *p*=0.005 and ENST00000518812, nominal *p*=0.03), both of which were upregulated with the risk allele. Note that ENST00000518812 is not a protein coding transcript.

## 4. DISCUSSION

In the last decade, GWASs have successfully identified relatively common genetic variants that drive disease associations, which has inspired a new chapter in genetic disease research that aims to identify the biological mechanisms through which genetic variation influences disease risk. Cellular modeling is quickly becoming a powerful tool for the study of genetic risk variants, particularly those within non-coding regions of the genome, as technological advances are being made in genome editing, iPSC generation from somatic cells, and directed differentiation methods^22^. Now, we are able to generate, study, and compare human neuronal cells with only a single nucleotide difference across the entire genome, as we and others have previously reported^21, 29, 66–70^. This approach provides the opportunity to elucidate the individual contributions of risk variants in a human- and disease-relevant system. Here, we apply this strategy to study a GWAS-identified, non-coding variant implicated in SCZ risk, rs4129585.

We first tested the regulatory potential of this variant in zebrafish and found that it drives reporter gene expression exclusively in the brain, and in regions relevant to SCZ, including areas involved in fear, anxiety, audio-sensory functions, and sensory-motor functions. Further, according to ENCODE data, this non-coding variant lies within a REST binding site, a TF already implicated in SCZ^44^. We next introduced the risk and non-risk alleles into iPSCs, differentiated them into biologically relevant NPCs, and produced clones homozygous for each allele on otherwise identical genomic backgrounds. Comparing the risk and reference alleles, we observed changes in the abundance of an alternatively spliced nearby gene, *ADGRB1,* as well as transcriptome changes affecting genes involved in axon guidance, synaptogenesis, and other functional pathways. The identified DE genes encode proteins that interact with each other more than would be expected by chance; further validating our results and supporting the observed functional enrichments.

Although we are not the first group to link synaptogenesis, axon guidance, and neuronal development with SCZ, we do so here for a specific SCZ risk variant, which we implicate as a regulator of a specific gene, *ADGRB1*, an adhesion G protein-coupled receptor^71^. Consistent with our transcriptomic observations, *ADGRB1* has been linked to brain development and developmental disorders^72, 73^, dendritic arborization^74^, synapse development and synaptic plasticity^65, 75, 76^. As such, it is not surprising to find suggestive evidence that it drives the risk in genetic associations with SCZ. The neighboring *TSNARE1* is also a strong candidate for SCZ risk, and it was originally reported as the driver of the association due to the localization of the lead variants within the gene^77^ in addition to its role in regulating endosomal trafficking in cortical neurons^78^.

The SNV rs4129585 has been previously identified as influencing the expression of nearby genes; in fact, GTEx shows that this variant is an eQTL for both *TSNARE1* and *ADGRB1.* However, since GTEx data is purely statistical – not functional – and comes from multiple tissues with bulk cells containing a variety of cell types, it only reflects the cumulative effects of all variants in LD at the locus where rs4129585 resides. There is no reason to assume that rs4129585 is the only functional variant at this locus, especially since there are other variants in LD with functional potential, such as rs13262595, as discussed in our methods. Our data shows that *ADGRB1* transcription is influenced by rs4129585 and by the sets of DE genes across the transcriptome, supporting the involvement of *ADGRB1* rather than *TSNARE1*, it remains possible that *TSNARE1* is also involved, but is regulated by another SNV in LD, such as rs13262595. Study of all such variants individually will shed additional light onto how these alleles and corresponding haplotypes influence SCZ risk and will provide valuable insights for the study of GWAS-identified variation, where the identification of multiple variants in strong LD is the rule, rather than the exception. Such future studies will eventually solve the puzzle of the complex interplay between multiple variants that can influence disease risk.

Our work suggests that, despite having a small effect on risk, a common variant can have a measurable effect on the transcriptome of NPCs, which points to specific functional pathways that, in turn, indicate disease mechanisms. There are three main reasons why this could be the case. First, studying risk and non-risk alleles in homozygosity, in otherwise isogenic cells, and under well-controlled conditions in vitro, may reduce noise and increase power to detect even small effects. Second, the study of cells in vitro not only reduces noise, but may also avoid a feedback buffering mechanism that acts at the organism level to dampen the effect of the variant^79^. Finally, studying an allele in isolation from other potentially functional alleles in LD might allow for the observation of effects that would otherwise be counteracted by the other alleles on the same haplotype. Natural selection could be making such a phenomenon more frequent than we recognize, if deleterious haplotypes are being selected against. This is particularly likely in SCZ, where negative selection appears to be strong^80–83^.

While studying NPC cultures in isolation has advantages, this isolation is also a limitation, since only cell-autonomous effects can be revealed. To bypass this limitation, 3D cerebral organoid culture systems could be leveraged, and accompanied by single cell sequencing. Despite being a more time consuming and expensive approach, it would be a valuable addition to monocellular cultures. While we do not currently have such results in organoid-based cultures or involving other GWAS-identified risk variants, we consider this, along with the study of multiple variants per locus, as an important addition to future projects.

## AUTHOR CONTRIBUTIONS

D.A. conceptualized the study design. R.J.B. designed and performed transgenic zebrafish reporter assays. M.H.W., C.Y., and B.R. performed cell culturing. M.H.W. performed genome editing of iPSCs, differentiation, and validation of NPCs. R.J.B. analyzed RT-qPCR data. D.A. performed bioinformatics analyses with contributions by R.J.B. R.J.B, D.A., and M.H.W. wrote the paper. M.H.W, R.J.B., C.Y., A.S.M., and D.A. contributed to the scientific discussion, data interpretation, and paper revision. All authors have read and agreed to the published version of the manuscript.

## FUNDING

This research was supported in part by awards from the National Institutes of Health (R01MH113215 and R21MH122936) to D.A., (R01HG010480 and R21NS128604) to A.S.M., and (T32GM007814-40) to R.J.B.; and by the Canadian Institutes of Health Research (DFD-181599) to R.J.B.

## DATA AVAILABILITY

RNA-sequencing data will be available at the Gene Expression Omnibus (GEO) upon article acceptance.

## ACKNOWLEDGEMENTS

The authors would like to acknowledge Anna Vakhnovetsky and Akul Umamageswaran for their technical support, as well as the FInZ Zebrafish Core Center for assistance with zebrafish husbandry.

## CONFLICTS OF INTEREST

The authors declare no competing interests.

## Notes

### Competing Interest Statement

The authors have declared no competing interest.

