## Supplemental Tables 1-4; Supplemental Figures 1-8 for "A Functional Schizophrenia-associated genetic variant near the *TSNARE1* and *ADGRB1* genes"

**Table S1.** Primers and candidate sequences used for Tol2-mediated transgenesis

| Name | Sequence (5' – 3') |
|---|---|
| Forward *attB* adapter + primer | GGGGACAAGTTTGTACAAAAAAGCAGGCT-GCCAAGAGTGAATCAGAC |
| Reverse *attB* adapter + primer | GGGGACCACTTTGTACAAGAAAGCTGGGT-ATCTCCTCTTCTGTAGACTG |
| Sequence containing **rs4129585**, (including primer annealing sites) | GCCAAGAGTGAATCAGACAAAGCCCTCAGCTTACTGGTTGGCGGGGGG<br>GCAGTCAACCCTCCTGGCAGGGACAGGGCTCTGCATCAGTGCACAGGA<br>CAAGGCGCACTGAATTAAGGACAACTTCAGGAC**A**CATGACAGCGCCCAT<br>GGCTTAGGATACTGGGGTTTGTGGGTGGCCAGTGTCTCTCAGGTCCATG<br>GTCCCCTCACCAGCATCGTCCCAGTCAGTAGTACAGCTGCCCTTTCCCA<br>AGCATCATCCCCCACCACGGCACAGAGGTGGGTCCCACCAACACCCCC<br>AGTCTACAGAAGAGGAGAT |

**Table S2.** Specific GFP expression patterns among founder zebrafish

| Location of GFP expression | Fraction of founder fish with expression (%) |
|---|---|
| Habenula | 4/4 (100%) |
| Caudal Hypothalamus | 4/4 (100%) |
| Inferior Olive | 4/4 (100%) |
| Glutamatergic neurons | 4/4 (100%) |
| GABAergic neurons | 4/4 (100%) |

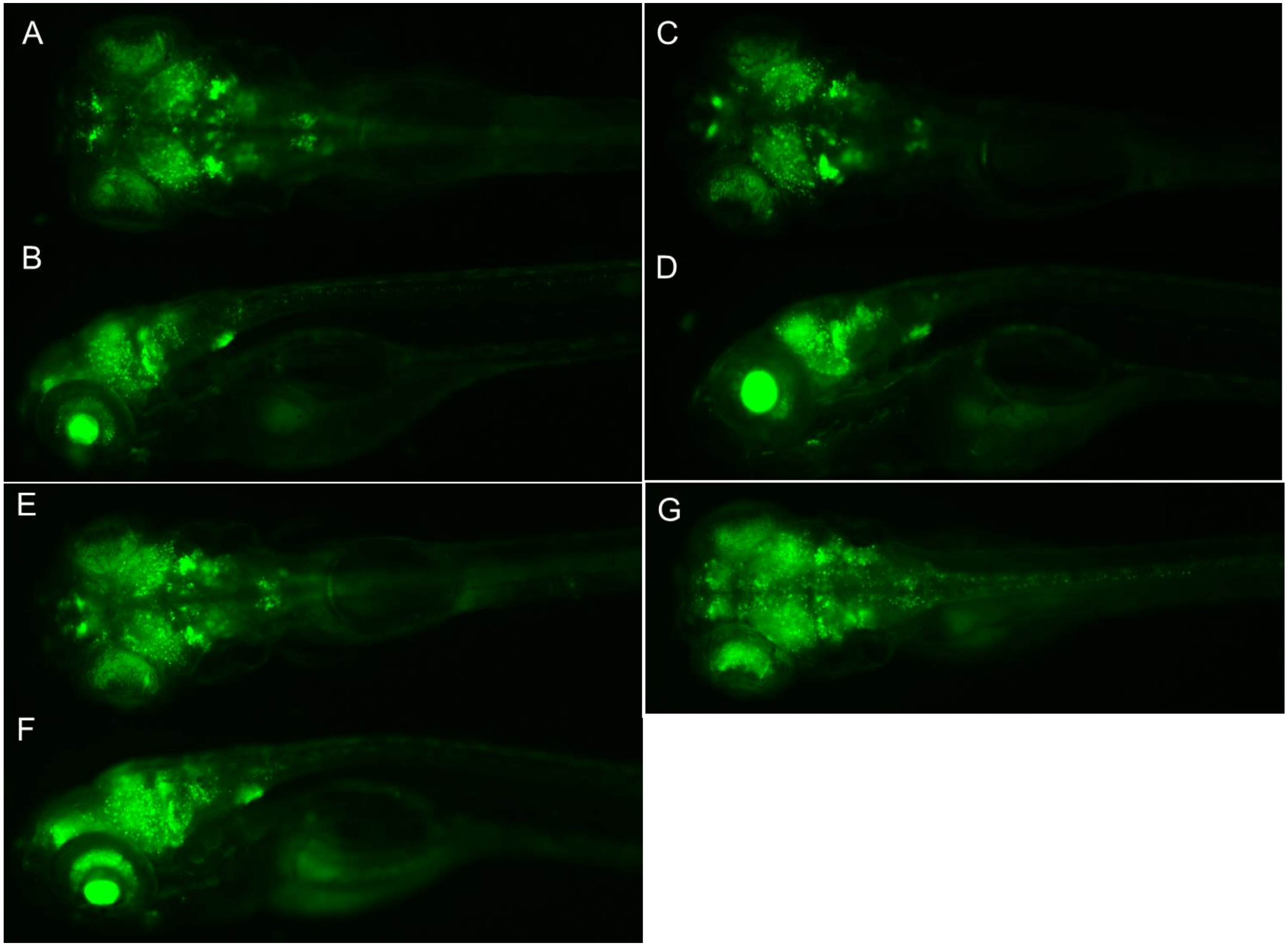


**Figure S1.** GFP expression patterns in the habenula, hypothalamus, inferior olive, and glutamatergic neurotransmitters in $F_2$ zebrafish larvae descending from 4 independent $F_1$ founders. **(A)** Dorsal and **(B)** lateral views of the founder zebrafish from Figure 1. Representative **(C)** dorsal and **(D)** lateral views of an $F_2$ offspring from founder zebrafish 2. Representative **(E)** dorsal and **(F)** lateral views of an $F_2$ offspring from founder zebrafish 3. A representative **(G)** dorsal view of an $F_2$ offspring from founder zebrafish 4.

**Table S3.** Primers for RT-qPCR

| Gene | Forward Primer (5' – 3') | Reverse Primer (5' – 3') | Category |
|---|---|---|---|
| *NES* | GAGGACCAGAGTATTGTGAGAC | CACAGTGGTGCTTGAGTTTC | NPC Marker |
| *PAX6* | TCTTTGCTTGGGAAATCCG | CTGCCCGTTCAACATCCTTAG | NPC Marker |
| *EMX1* | CGAGACGCAGGTGAAGGTG | CTGCCCTCGTGGGTTTGT | NPC Marker |
| *FOXG1* | GCAGCACTTTGAGTTACAACGGCA | AGTTCTGAGTCAACACGGAGCTGT | NPC Marker |
| *OCT4* | AGTTTGTGCCAGGGTTTTTG | ACTTCACCTTCCCTCCAACC | iPSC Marker |
| *NANOG* | ACAACTGGCCGAAGAATAGCA | GGTTCCCAGTCGGGTTCAC | iPSC Marker |
| *TRA-160* | ACAGGAAACACCCTCTGTGC | GAAGGTGGCTTTGACTGCTC | iPSC Marker |
| *GAPDH* | CAACTACATGGTTTACATGTTCCAA | GACAAGCTTCCCGTTCTCAG | Loading Control |
| *ACTB* | GTCTTCCCCTCCATCGTG | AGGATGCCTCTCTTGCTCTG | Loading Control |

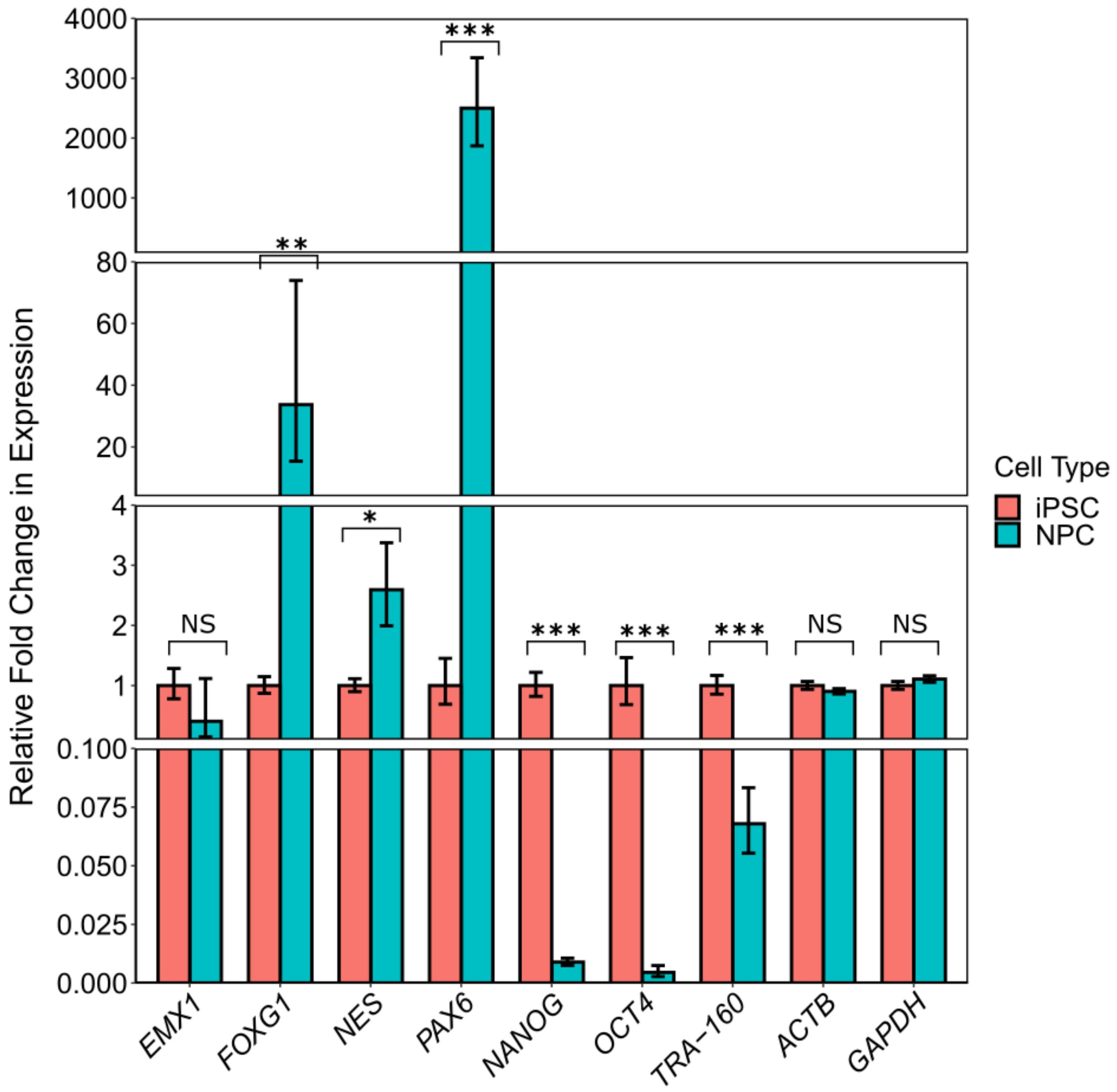


**Figure S2.** Successful differentiation from iPSCs to NPCs is confirmed by expression of iPSC and NPC markers by RT-qPCR. Relative to normalized gene expression in iPSCs, expression of iPSC marker genes (*NANOG, OCT4*, and *TRA-160*) diminishes in differentiated NPC clones. Expression of NPC marker genes (*FOXG1, NES, PAX6*) increases in NPCs, with the exception of *EMX1*, which did not exhibit significant fold-change in expression. As expected, controls *ACTB* and *GAPHD* did not exhibit significant fold-change in expression between iPSCs and NPCs. Data represents 2-3 technical replicates from each of 7 biological replicates (REF_1, REF_4, REF_6, RSK_1, RSK_2, RSK_3, RSK_4). NS = not significant; * = $p$ < 0.01; ** = $p$ < 0.001; *** = $p$ < 0.0001.

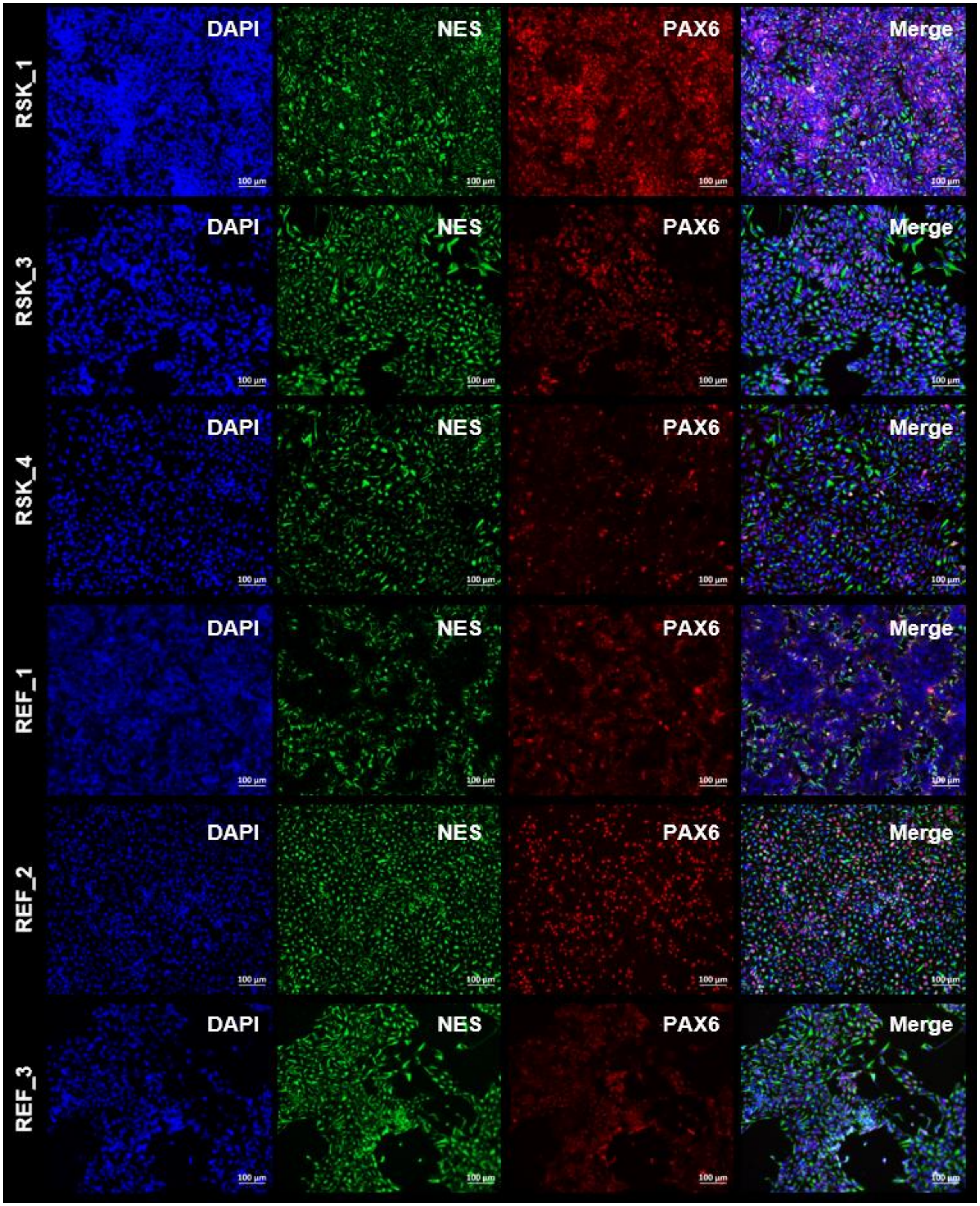


**Figure S3.** Representative immunocytochemistry staining with DAPI, NES, and PAX6 confirms successful differentiation of RSK and REF clones into NPCs.

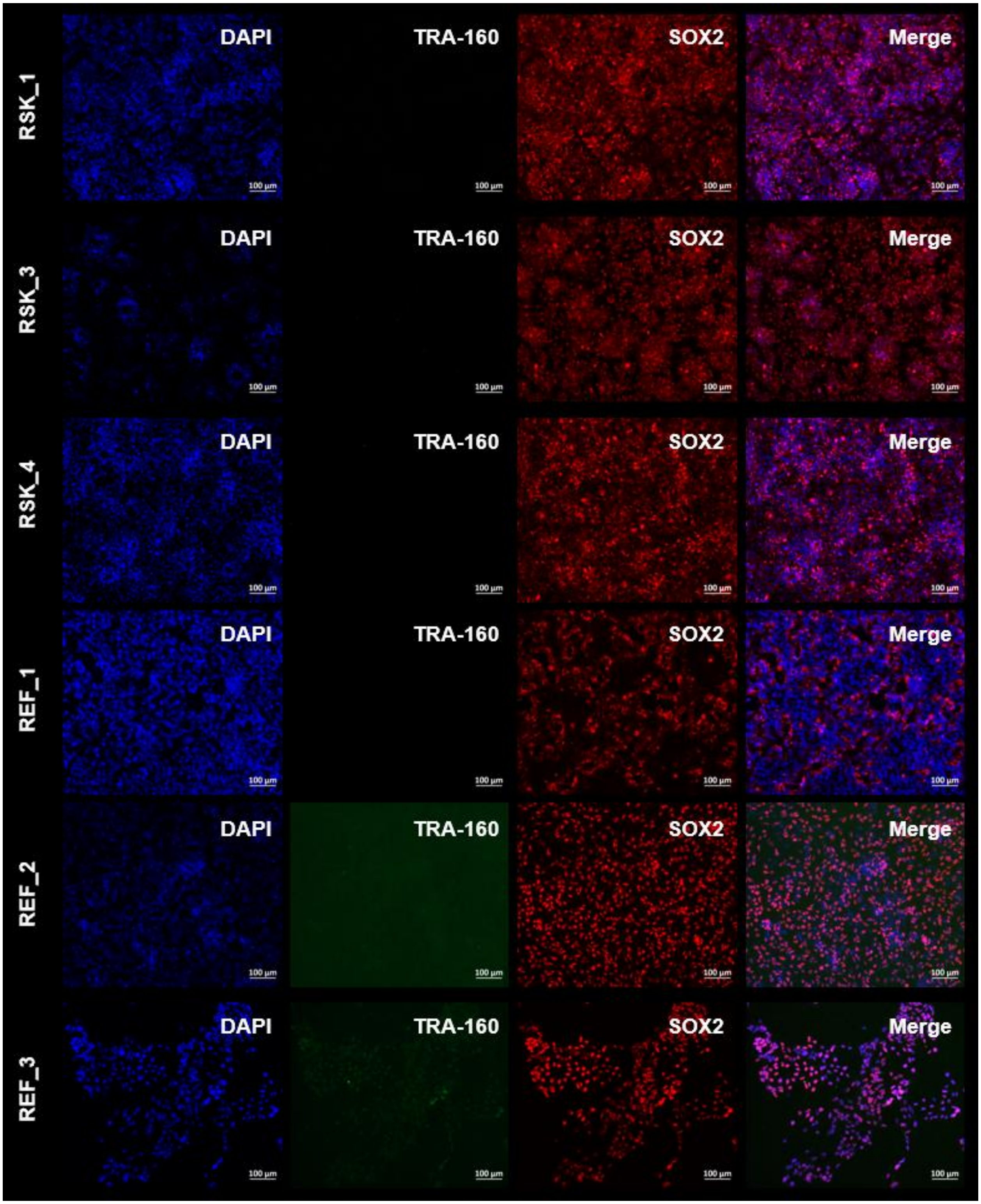


**Figure S4.** Representative immunocytochemistry staining with DAPI, TRA-160, and SOX2 confirms successful differentiation from iPSCs into NPCs.

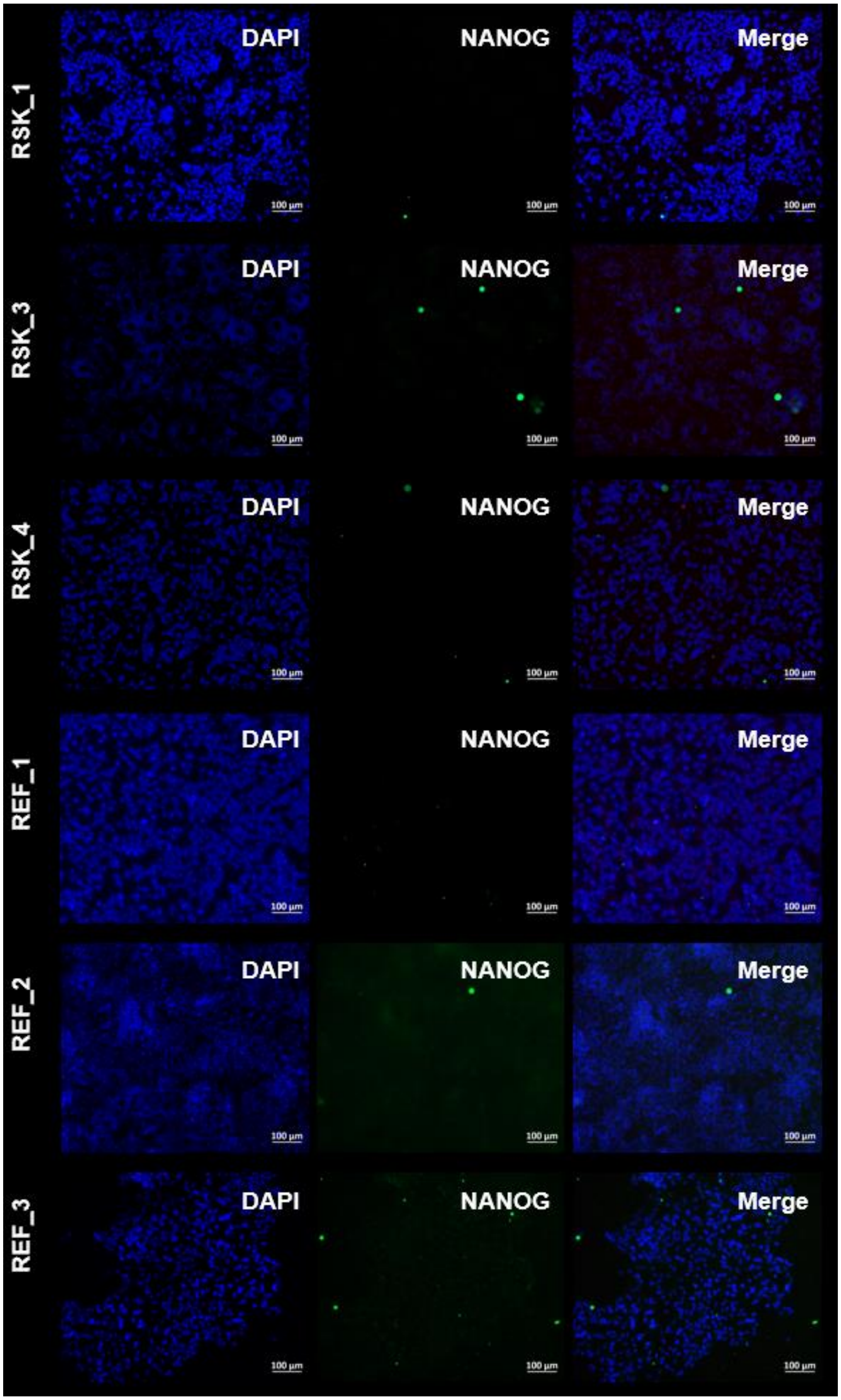


**Figure S5.** Representative immunocytochemistry staining with DAPI and NANOG confirms that the differentiated cells are no longer iPSCs.

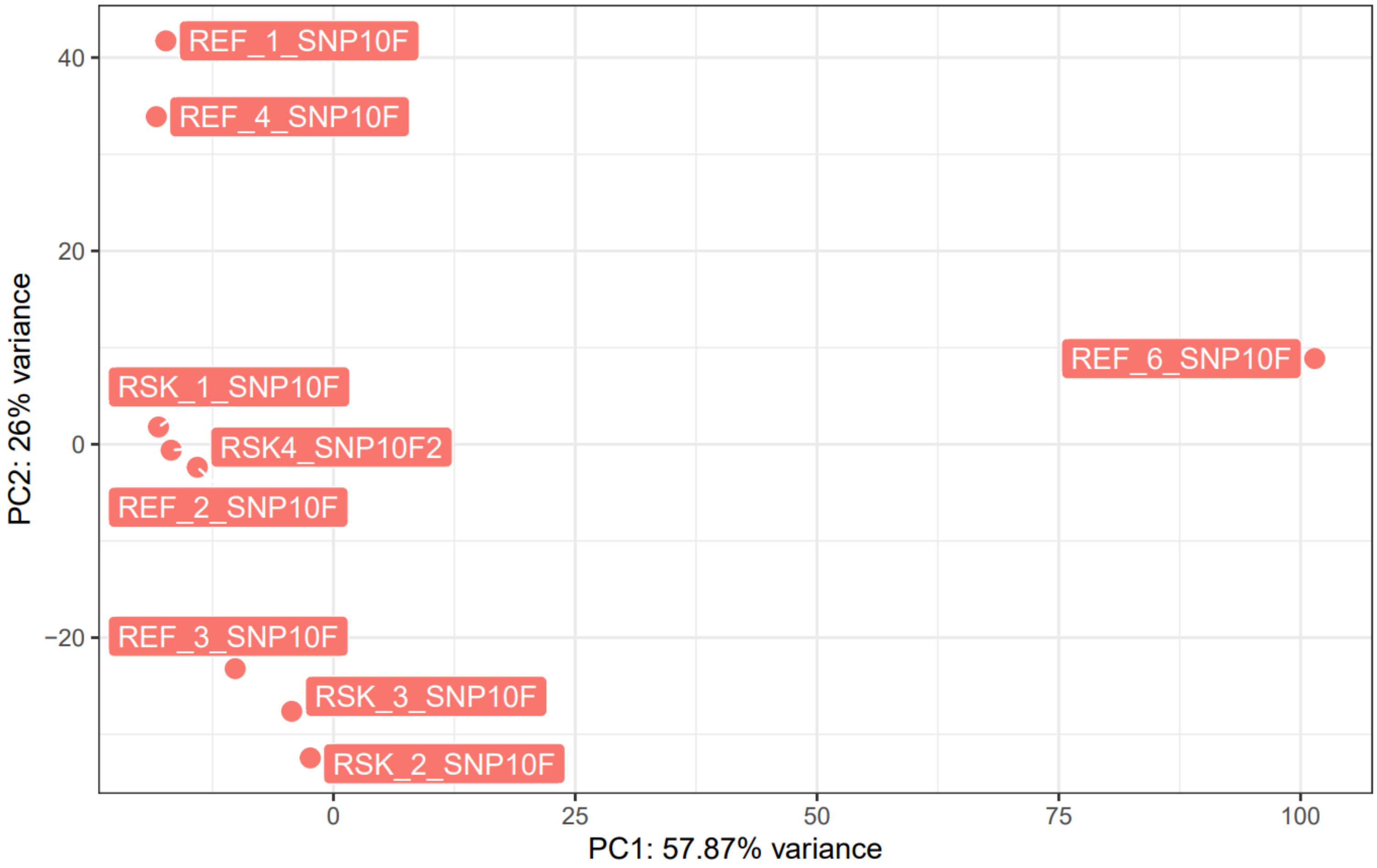


**Figure S6.** PCA plot of RNA-seq data shows that one iPSC-derived NPC clone carrying the reference allele (REF_6) is an outlier across PC1, but that all other samples cluster by their status as carrying the reference (REF) or risk (RSK) allele.

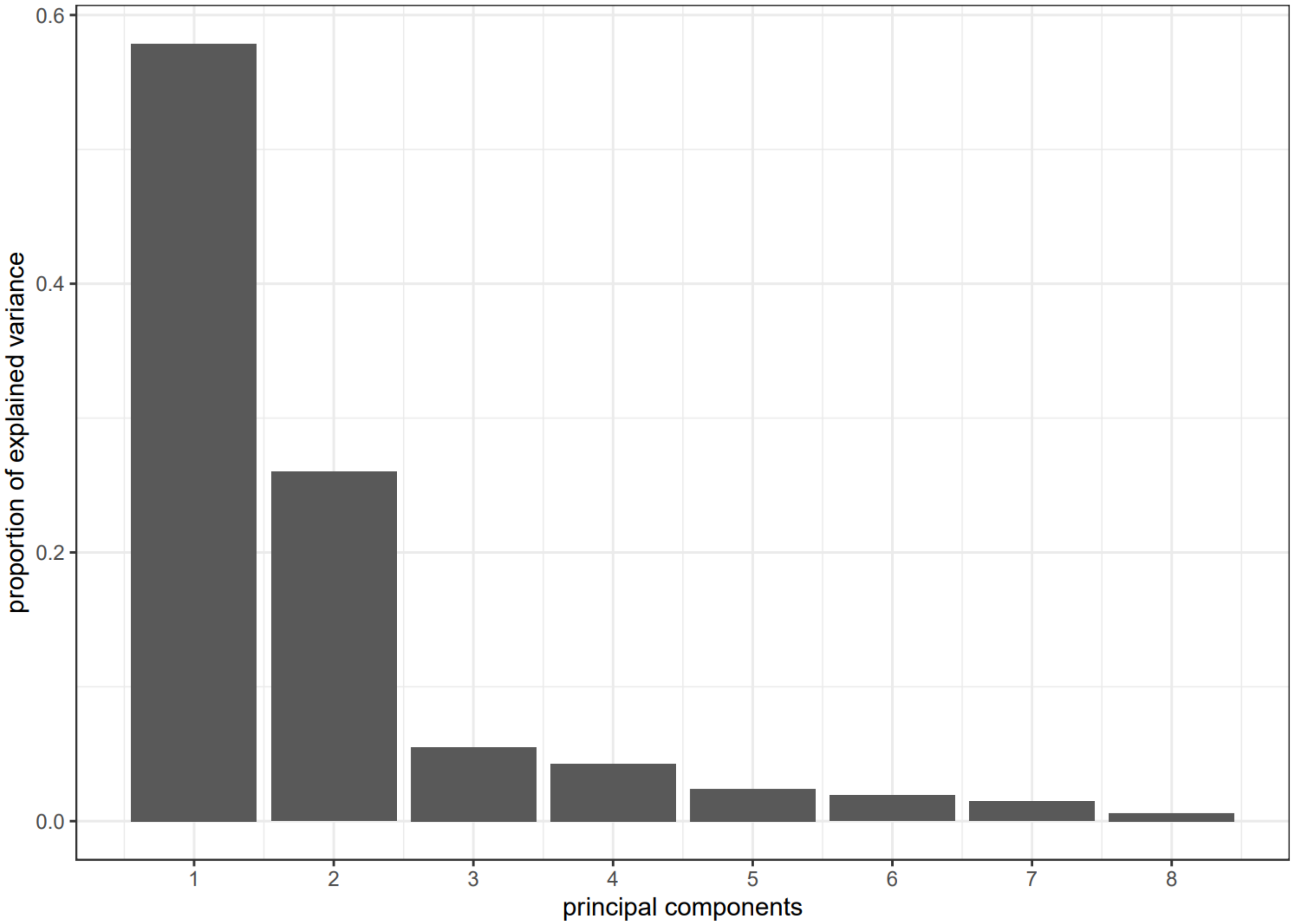


**Figure S7.** Scree plot of RNA-seq PCA shows that the majority of the sample variance is explained by PC1 and PC2.

**Table S4.** Normalized read counts of marker genes from iPSC-derived NPCs containing the reference (REF) or risk allele (RSK).

| Gene | REF_1_S NP10F | REF_2_S NP10F | REF_3_S NP10F | REF_4_S NP10F | REF_6_S NP10F | RSK_1_S NP10F | RSK_2_S NP10F | RSK_3_S NP10F | RSK_4_S NP10F2 |
|---|---|---|---|---|---|---|---|---|---|
| *NES* | 65403 | 42561 | 38041 | 60266 | 15405 | 22256 | 48715 | 14428 | 18165 |
| *PAX6* | 1933 | 5241 | 10154 | 5305 | 0 | 10162 | 10083 | 8781 | 10035 |
| *EMX1* | 6 | 4 | 888 | 4 | 58 | 224 | 922 | 504 | 198 |
| *FOXG1* | 36 | 414 | 6499 | 82 | 8 | 918 | 12853 | 2481 | 1037 |
| *OCT4 (POU5F1)* | 1 | 6 | 3 | 0 | 78437 | 4 | 11 | 5 | 0 |
| *NANOG* | 7 | 2 | 6 | 22 | 2463 | 24 | 30 | 34 | 14 |
| *TRA-160 (PODXL)* | 3796 | 12402 | 5298 | 6442 | 101164 | 6711 | 2813 | 2142 | 5972 |

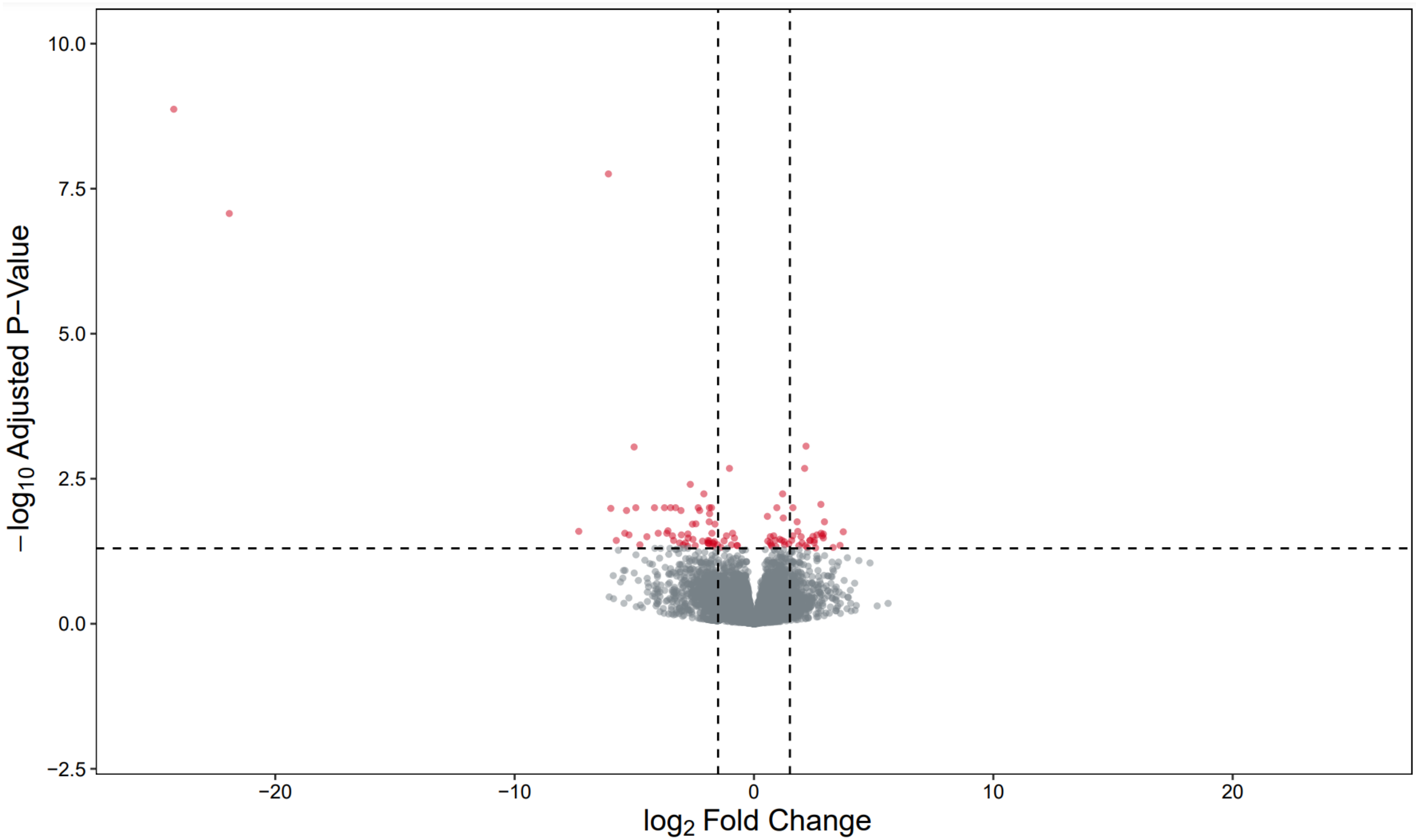


**Figure S8.** Volcano plot of $-\log_{10}$ adjusted p-value versus $\log_2$ fold change with DESeq2, contrasting the fold change in expression of NPCs containing the risk allele, using NPCs containing the reference allele as reference.
